# Slow and steady: universal learnable structure in animal vocalizations

**DOI:** 10.1101/2024.06.20.599962

**Authors:** Ronald W. Di Tullio, Linran Wei, Vijay Balasubramanian

## Abstract

The discovery of universal structures in natural stimuli has allowed neuroscientists to make normative statements about the organization of sensory systems. We sought such structure in animal vocalizations – stereotyped sound patterns that are received, recognized, and responded to by conspecific and allospecific individuals. Across humans, macaques, and four evolutionarily divergent species of birds we found that: (a) Vocalizations have the majority of their power in low-frequency temporal modulations, (b) Most of this power arises from 9-10 “slow” auditory features, (c) The acoustic space spanned by the leading slow features is universal,up to a small rotation, across species, (d) The leading slow features span the same space as the features most predictive of the future time course, (e) A classifier can learn to discriminate vocalizations with high accuracy based on the 10 leading features. These results suggest a universal temporal structure in vocalizations that may be exploited in animal communication.

## I. INTRODUCTION

All organisms exploit patterns in the world to make inferences and select actions. Species relying on acoustic communication use patterns in sound to discriminate both conspecific and allospecific vocalizations as well as guide behavior[1–10]. Here, we find a universal, learnable, low-dimensional structure in the vocalizations of two primate and four evolutionarily distinct bird species. Our results suggest that the acoustic structure we reveal may be shared across species that use a vocal apparatus to transmit messages.

Vocalizations are vibrations produced by specialized organs in conjunction with the respiratory system. These vibrations propagate through the surrounding medium and are received and processed by the auditory systems of other individuals [2, 6, 11]. Vocalizations can vary significantly across species, and even between individuals of the same species, but nevertheless can show striking regularities. Notably, there is a conserved tempo in isochronous animal communication [12, 13]. If richer regularities are present, they could indicate a universal structure that allows animals to learn, discriminate, and parse communications from conspecific and, to a varying extent, allospecific individuals [1, 3, 14–17]. Thus, we looked for, and sought to characterize, shared structure in vocalizations of two primate species (humans, macaques) and four bird species (American yellow warbler, blue jay, great blue heron, and house finch), selected for the ready availability of well-recorded data.

We first analyzed modulation spectra of vocalizations. Expanding on previous observations [18, 19], we found that, across species, vocalizations concentrate power at lower temporal modulations. We then showed that the structure arising from this concentration of power can be learned by Slow Feature Analysis (SFA), an unsupervised learning algorithm that learns the most temporally regular (colloquially “slowest”) parts of a signal [20–24]. Given this correspondence between low temporal modulation and “slowness” as defined by SFA, we focused on analyzing the structure learned by SFA. We found that vocalizations across species contain approximately 9-11 slow SFA features that define a low dimensional subspace of acoustic space.

We further found that, within species, the low dimensional subspaces characterizing particular vocalizations can be minimally rotated (20 degrees or less) to overlap, indicating shared temporal structure defined by the slowest SFA features. Surprisingly, we found that the same was true for the acoustic subspaces characterizing different species: the subspaces defined by the slowest SFA features can be minimally rotated (20 degrees or less) to overlap. This suggests a universal learnable temporal structure inherent to animal vocalizations that can be discovered by a simple, unsupervised learning algorithm.

Animals sense the world in order to adapt their behavior to predicted future states. This suggests that sensory systems should be tuned specifically to *predictive* components of environmental signals. If so, we would expect auditory receivers and pathways to extract and process the most predictable components of vocalizations. We extracted these using Predictable Feature Analysis (PFA) [25, 26], and found that the leading 5 PFA features span the same acoustic space as the leading SFA features. Thus learning the “slowest” features of vocalizations also learns their most predictable features. Note that “fast features” could also be predictable provided they contained the requisite temporal correlations – apparently, they do not.

Finally, we tested whether the universal acoustic subspace learned by SFA supports a core auditory task: discriminating vocalizations. To this end, we trained a network to classify vocalization pairs that had been projected into the universal subspace. We found: 1) classification performance was high (*>* 97%), 2) a network using only the 10 slowest features had roughly equivalent performance to a network using twice as many features, 3) network performance depended most on the slowest features and drops precipitously if these are shuffled. These results show that the universal acoustic subspace learned by SFA can be exploited to solve auditory tasks.

Our findings suggest that, despite apparent spectral differences, vocalizations of different individuals and species share a universal temporal structure that could be, and likely is, exploited in animal communication.

## II. RESULTS

### A. Temporal regularity in vocalizations

Auditory objects, recurring acoustic patterns identifiable as “a sound”, are defined by regularities in their spectro-temporal dynamics. We thus looked for structure in these dynamics that could be exploited by a learner. Previous work suggests several temporal regularities [18, 19, 27–31], including coincidence, correlation, and continuity of acoustic features (Fig. 1A). Thus, we sought to characterize such regularities in the vocalizations of several species – birds (great blue heron, house finch, American yellow warbler, and blue jay), humans (English speakers saying digits 0-5), and macaques (producing *coo* vocalizations) (Methods - Auditory Stimuli).

**FIG. 1.**
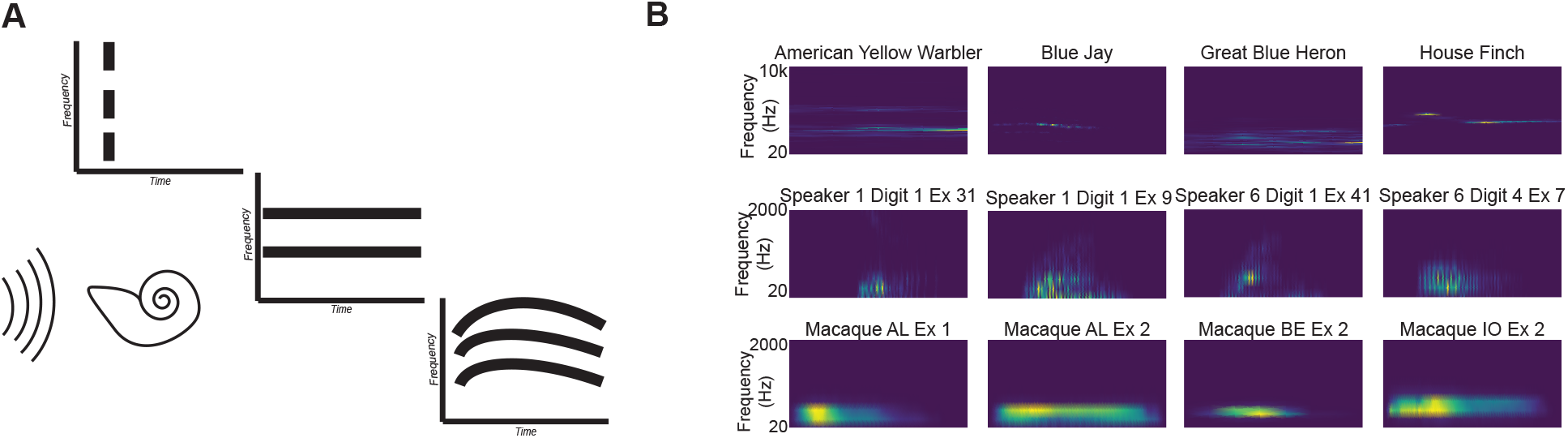
Coincidence and continuity – vocalization spectrograms. **A**. *Left*. Schematic of auditory stimuli received by the auditory system (cartoon cochlea). *Middle*. Coincident onset *Right*. Coincident onset and correlation. *Far Right*. Continuous and consistent varying correlations between the frequency components of an auditory stimulus. **B**. Example spectrograms for birds, humans, and macaques. Warmer colors indicate more energy at that frequency-time point and cooler colors, less. Spectrograms temporal scale varies for clarity(durations are *∼* 50 ms for birds and up to 500 ms for macaques). *Top*. Calls from four bird species. *Middle*. English speakers pronouncing digits 0-5. *Bottom*. Macaques producing *coo* vocalizations. Different examples (Ex) of the same vocalization by a given speaker shown for humans and macaques.

We analyzed the acoustic structure of vocalizations via their spectrograms: the Fourier power spectra taken in small temporal windows [18, 19, 31–34] (Fig. 1B). Qualitatively, vocalizations produced by each species comprise different sets of frequencies, but all manifest correlations that change in a regular manner over time. We quantified this temporal structure by averaging the modulation power spectrum (MPS) for vocalizations within each animal group (Fig. 2A). We defined the MPS as the 2d Fourier power spectrum of a cochleagram; a spectrogram generated from a model of cochlear processing (Methods - Simple Cochlear Model). The MPS displays acoustic energy in temporal and spectral frequency bands.

**FIG. 2.**
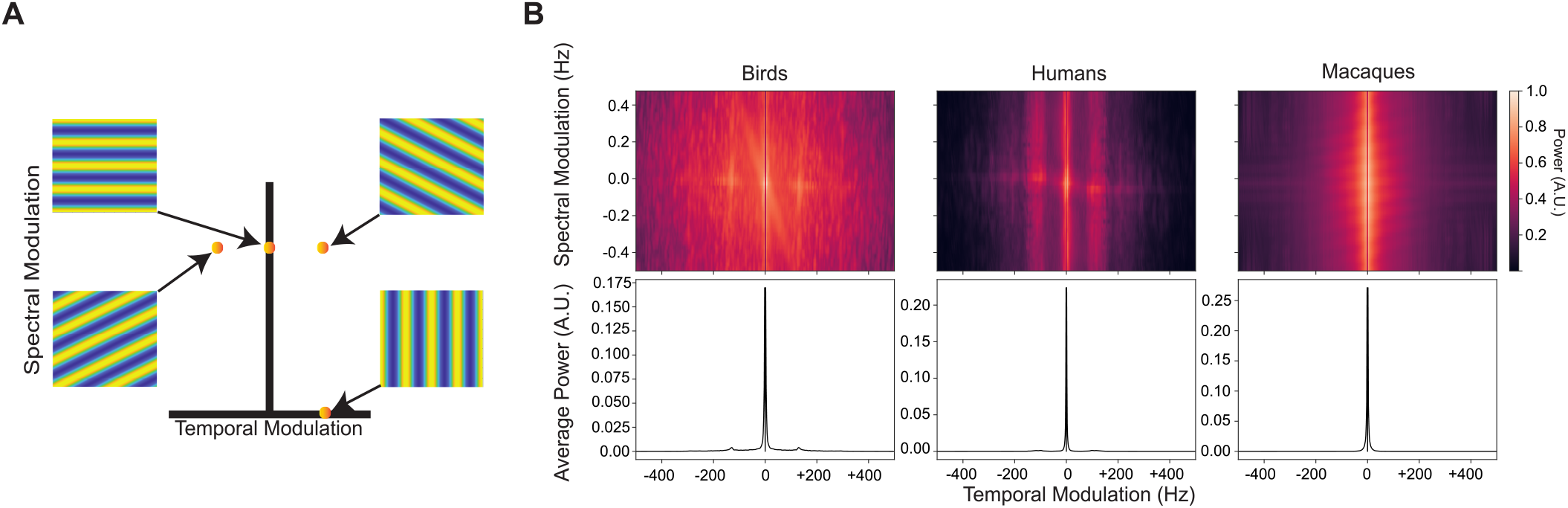
Coincidence and continuity – modulation spectra. **A**. A spectrogram or cochleagram can be decomposed as a weighted sum of spectral- and temporal-modulation rates. Upper left - spectrogram with only spectral modulation Lower right - a spectrogram with only temporal modulation. The other two spectrograms are examples with both spectral- and temporal-modulation. Color represents energy at each time-frequency point. **B (Upper)**. Average spectrotemporal modulation spectra for each animal group. **B (Lower)**. Average spectral power as a function of temporal modulation. Modulation spectra are calculated following [18, 19]. 200 vocalizations were used to generate each plot. These vocalizations comprised the pairs were used in all subsequent analyses and figures unless stated otherwise (Methods - Auditory Stimuli).

We found that our vocalizations consisted primarily of components with low temporal modulation (Fig. 2B, upper panel). Averaging across spectral modulation for each group makes this concentration at lower temporal modulations clearer (Fig. 2B, lower panel). This finding replicates and expands upon [18, 19] suggesting that animal vocalizations across species are dominated by temporally regular acoustic features.

### B. Slow features summarize temporal regularity in vocalizations

The concentration of power at low temporal modulations in the MPS suggested that an agent learning vocalizations could focus attention on acoustic features that change relatively slowly. We have previously shown that Slow Feature Analysis (SFA) [20–23, 35–37] extracts such features from macaque vocalizations [31]. Briefly, SFA extracts from a signal the features that have the lowest temporal variance and thus change “slowly” over time (Methods - Slow Feature Analysis).

To verify that features learned by SFA capture the regularities responsible for the power in the low temporal modulation, we analyzed our vocalizations with SFA and projected onto the 10 slowest features. We then took these projections and repeated our MPS analysis. If SFA is capturing temporal regularity in the manner we are proposing, then the MPS for the slowest 10 features should capture the large peak seen at slower temporal modulation frequencies for the complete signal. Indeed, this is what we find (Fig. 3). These results indicate that learning slow features via SFA successfully captures the structure we observed in natural animal vocalizations.

**FIG. 3.**
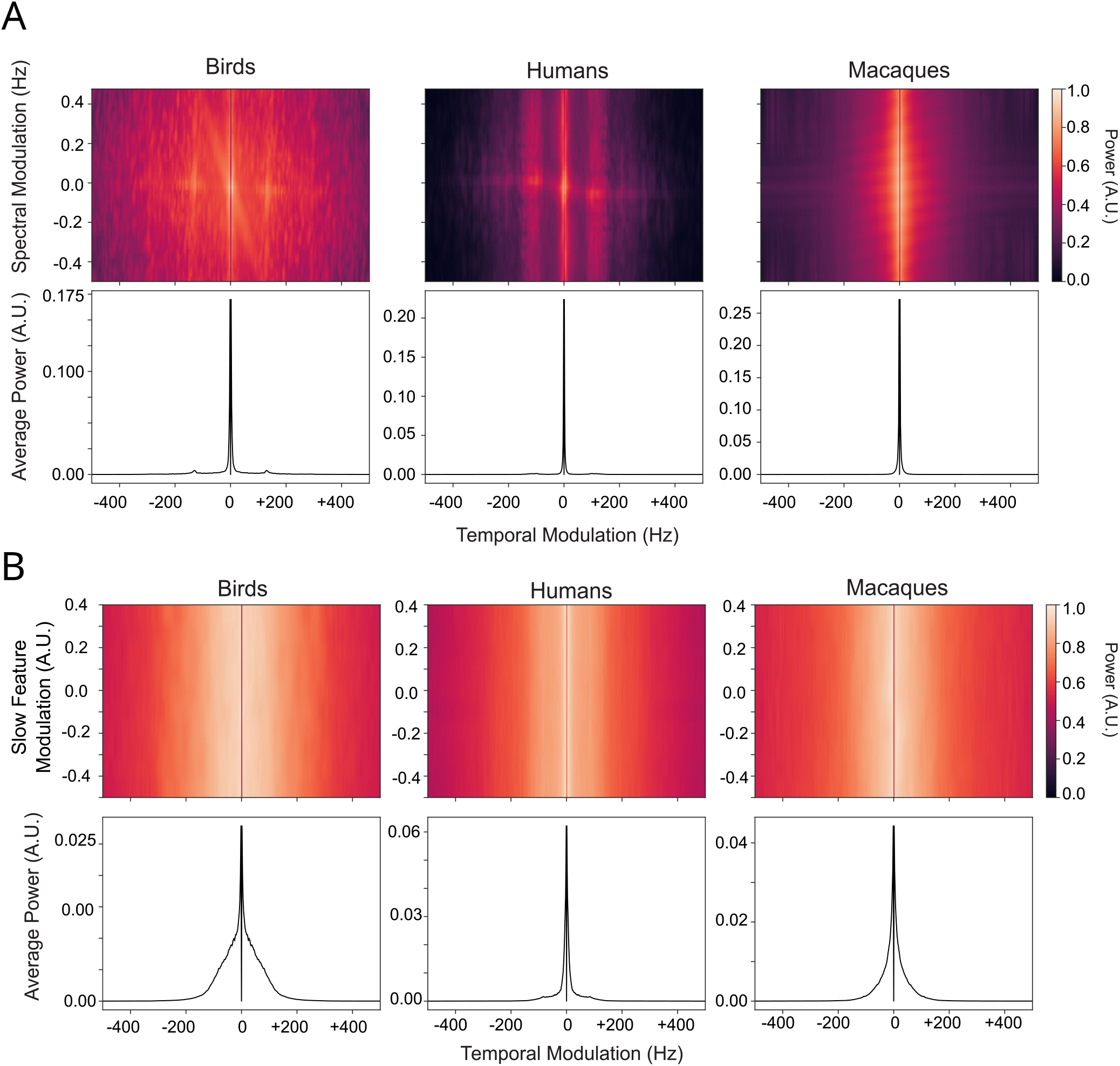
Modulation power spectrum reconstruction with the 10 slowest SFA features. As with previous plot, the heatmap is the average modulation power spectrum (MPS) with the average taken across our 100 example vocalizations. The line plot below is the average power in that temporal modultion bin with the average take across all spectral modulation bins. **A**. The original MPS with the 0 Hz temporal modulation (DC component) removed. **B**. Reconstructing the MPS using the 10 slowest SFA features.

### C. Vocalizations consist of few, very slow features

Previous studies have suggested how SFA or SFA-like computations could be performed by neural circuits [21, 31, 37]. Furthermore, the MPS reconstruction results from above suggest that a relatively small number of slow features should be sufficient to capture the temporally regular components of acoustic structure. To quantify this, we used the fact that we formulated SFA as a generalized eigenvalue problem (see Methods) and directly analyzed the eigenvalue distribution. We extracted this distribution for each vocalization and averaged within each animal group. Our MPS reconstruction suggests that there should be an inflection point, i.e., a relatively sharp slope change, when we plot the eigenvalue distribution as a function of eigenvector index. Fig. 4 indeed shows a decrease in the slope of the eigenvalue distribution for all animal groups at around the tenth slowest feature. An inflection point analysis using the KNEEDLE algorithm [38] confirms this observation for each animal group (Birds: 11th, Humans: 9th, Macaques: 8th; colored triangles in Fig. 4).

**FIG. 4.**
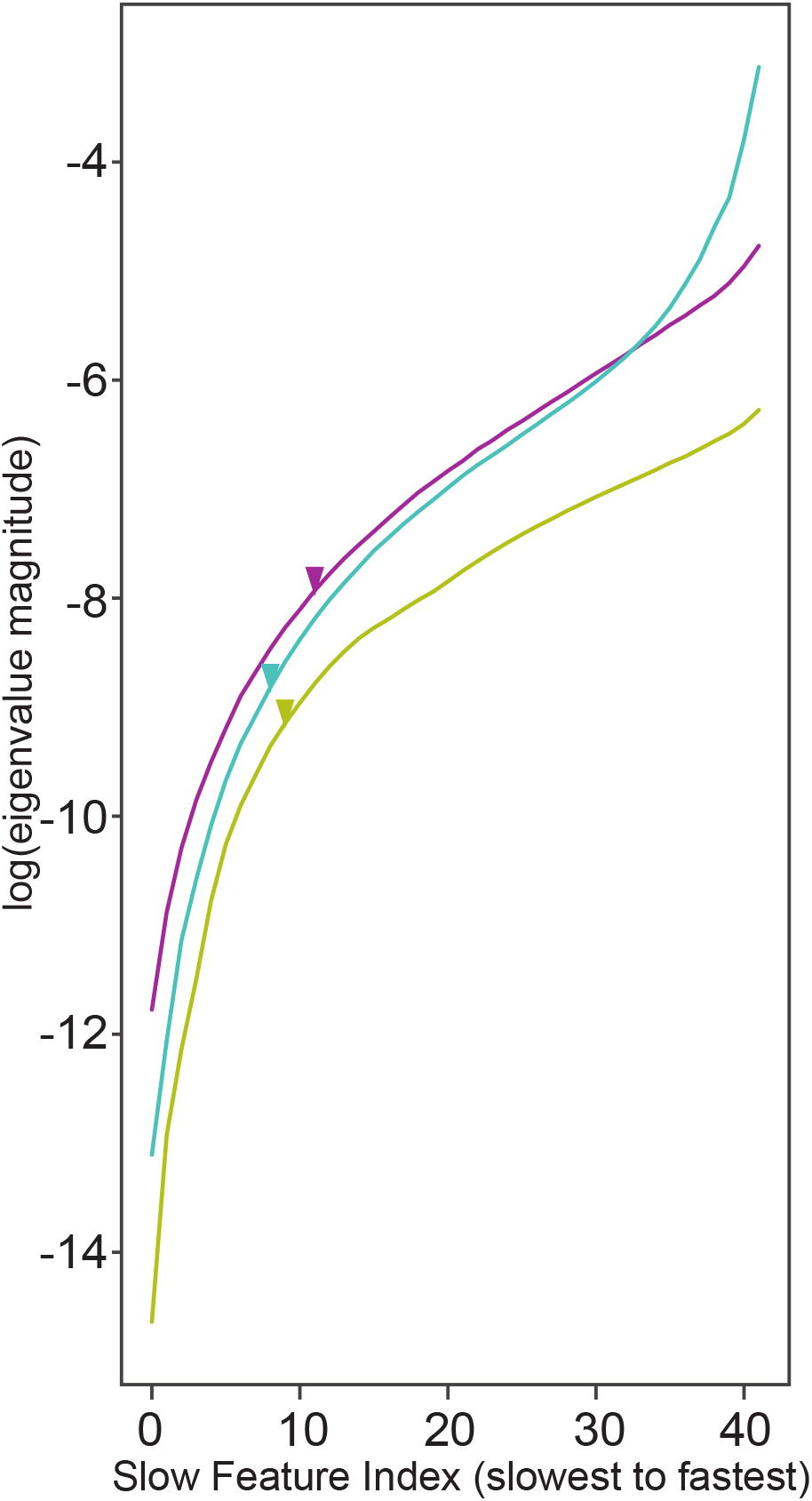
Slowness as a function of SFA eigen value index. Plot of the logarithm of the eigenvalues (y-axis) associated with each slow feature index (x-axis) for each animal group. A smaller eigenvalue indicates a slower feature. Magenta = birds; Yellow = humans; Cyan = macaques.

This analysis confirms that animal vocalizations are comprised of a small number of slow features and consequently suggests that an agent trying to exploit temporal regularities in vocalizations would not need to learn the full spectrum of slow features, but rather a limited subset. In other words, animal vocalizations reside in a relatively low dimensional acoustic subspace that can be found by SFA or SFA-like computations.

### D. The slow features of animal vocalizations define a shared subspace within animal groups

We next asked whether the low dimensional acoustic subspaces uncovered by SFA are shared across vocalizations and individuals within each group. To do so, we randomly paired vocalizations within animal groups (birds, humans, and macaques), extracted each of their slow features with SFA, and constructed an “SFA-Matrix” where each column is an SFA feature for each vocalization. We then calculated and analyzed the Gram matrix for each vocalization pair (Methods - The Gram Matrix and Subspace Overlap). In brief, the Gram matrix is the inner product between two matrices and can be analyzed to quantify the overlap between two subspaces. We then took the average Gram matrix for all pairs for each animal group. We found that, on average, the slowest 10 vocalization features show high overlap Fig. 5 A). In effect, the dominant acoustic subspaces associated with different vocalizations within an animal group are largely the same.

**FIG. 5.**
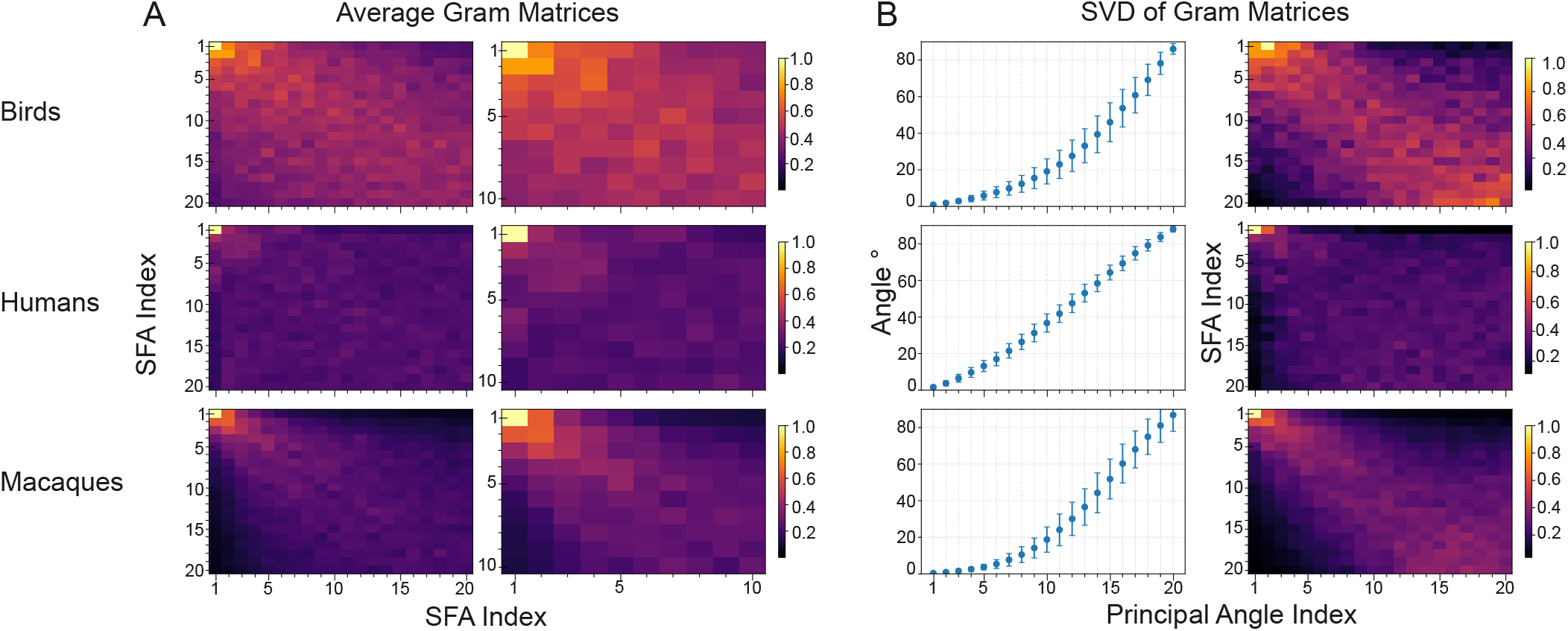
Slow Feature Subspaces within Animal Group. **A. Left Column**. Plot of the average gram matrix for 100 pairs of slow features for each animal group. Average is taken across pairs. Brighter spots indicate stronger overlap between those particular slow feature pairs (e.g. a bright spot in position 1,1 means the slowest feature, on average, overlaps strongly between pairs). **A. Right Column**. Zoom in to the 10 slowest features of the plot in the leftmost column for each animal group. **B. Left Column**. Average principal angles for the gram matrices of each animal group. Average is take across pairs. 0 degrees indicates a direction of perfect overlap, 90 degrees indicates orthogonality. **B. Right Column**. Average contribution of each slow feature to each principal angle for each animal group. Brighter spots indicate that features contributes more strongly to that principal angle and darker spots indicate little to no contribution.

The Gram matrix itself can be further analyzed via singular value decomposition (SVD). This analysis generates two important quantities–principal angles and the contribution of each slow feature to the principal angles. If the subspaces defined by our slowest SFA features overlap significantly, we would expect to see: 1) a subset of principal angles with magnitudes near zero 2) the slowest SFA features contribute the most to these low magnitude principal angles. We used ∼ 20 degrees as our cutoff for an angle being “near zero” and found that both conditions are satisfied (Fig. 5 B). For each animal group, there are several (approximately 5-9) principal angles that are less than 20 degrees and the slowest SFA features contribute the most to these. We also observe the converse: the faster features contribute the most to the orthogonal (90 degree) principal angles.

Taken together, these results suggest that there is a universal acoustic subspace of temporally regular signals within which vocalizations reside for each animal group.

### E. The slow features of animal vocalizations define a shared subspace between animal groups

If the structure we have found is universal, it should exist not just within but across animal groups. To test whether we have found such a structure, we repeated our Gram matrix analysis across animal groups (Fig. 6). We found that the slowest SFA features show the highest similarity across animal groups, with the highest concentration around the 5 slowest. Likewise, the principal angle distribution contains approximately 5-7 principal angles that are less than 20 degrees, and we find that the slowest features contribute the most to the overlapping angles, with the slowest 5 features showing the most contributions. We again find the expected converse–the fastest features contribute the most to the orthogonal angles.

**FIG. 6.**
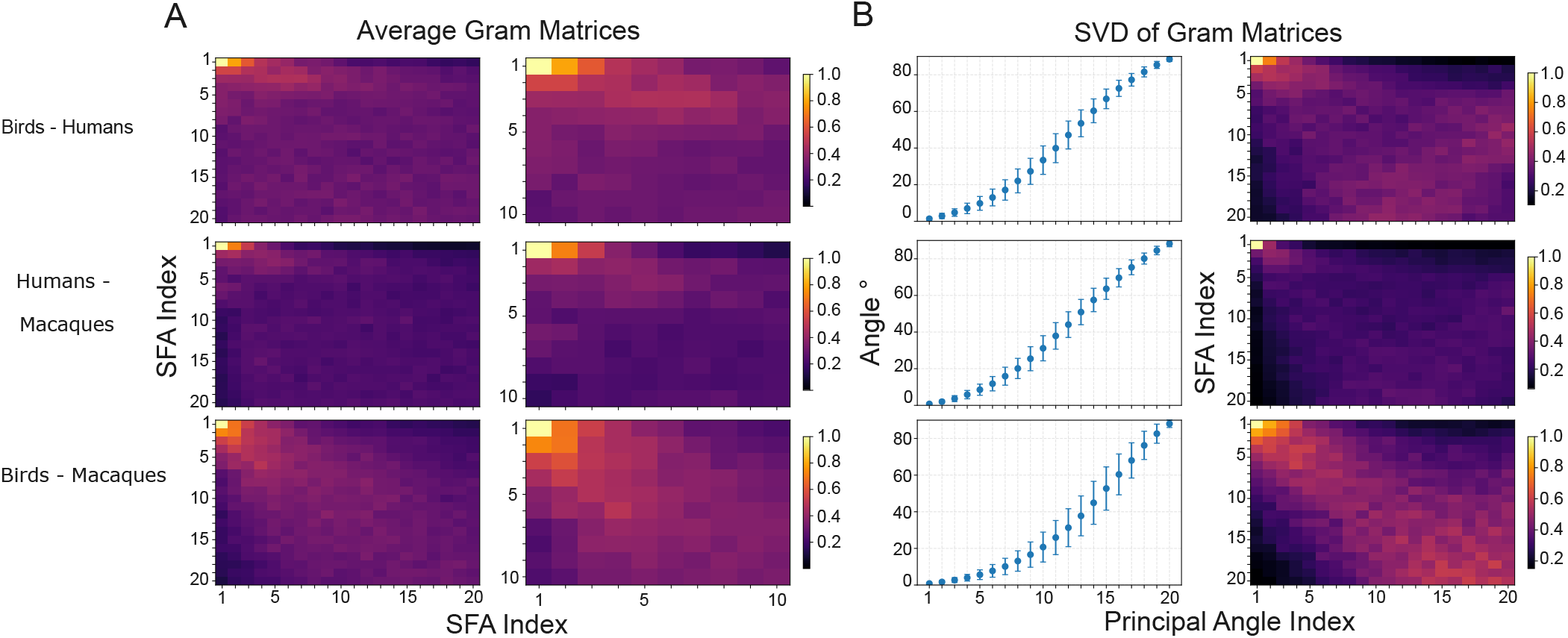
Slow Feature Subspaces between Animal Groups. **A. Left Column**. Plot of the average gram matrix for 100 pairs of slow features for each animal group pair. Average is taken across vocalization pairs. Brighter spots indicate stronger overlap between those particular slow feature pairs (e.g. a bright spot in position 1,1 means the slowest feature, on average, overlaps strongly between pairs). **A. Right Column**. Zoom in to the 10 slowest features of the plot in the leftmost column for each animal group pair. **B. Left Column**. Average principal angles for the gram matrices of each animal group pair. Average is take across vocalization pairs. 0 degrees indicates a direction of perfect overlap, 90 degrees indicates orthogonality. **B. Right Column**. Average contribution of each slow feature to each principal angle for each animal group pair. Average is take across vocalization pairs. Brighter spots indicate that features contributes more strongly to that principal angle and darker spots indicate little to no contribution.

These results indicate that despite the large variance in production mechanisms and spectral content, vocalizations from birds, humans, and macaques share underlying temporal regularities that are readily learned by algorithms like SFA, and constitute a low dimensional acoustic subspace defined by the approximately first 5 to 10 slowest SFA features. These results also coincide with our earlier findings: there was an inflection point in the slowness of the slow features around the 10th slowest feature and the shared subspaces we have found appear to be 10 dimensional or lower.

### F. Predictability equals slowness

We could have alternatively viewed temporal regularity through the lens of predictability. In such a framing, vocalizations are assumed to be autoregressive, i.e. one can predict what is coming next given solely what has happened so far. Note that predictability is not the same as continuity or smoothness, because highly discontinuous signals can be completely predictable from their pasts if they are generated deterministically. In the language of slowness, a fast but repetitive signal would still be predictable but will not be picked up in the slower features of SFA if there are parts of the signal that have lower temporal variance. We thus sought to determine the relationship between the predictable and slow features of vocalizations.

We decomposed individual vocalizations using Predictable Feature Analysis (PFA)[25, 26] as well as SFA, and used Canonical Correlation Analysis (CCA) [39–42] to compare PFA and SFA features. Briefly, CCA compares multidimensional data sets by finding projections maximizing correlation between the sets (Methods - PFA SFA Comparison). We did this for each vocalization separately and aggregated vocalizations by animal group: birds, humans, macaques. (Fig. 7)

**FIG. 7.**
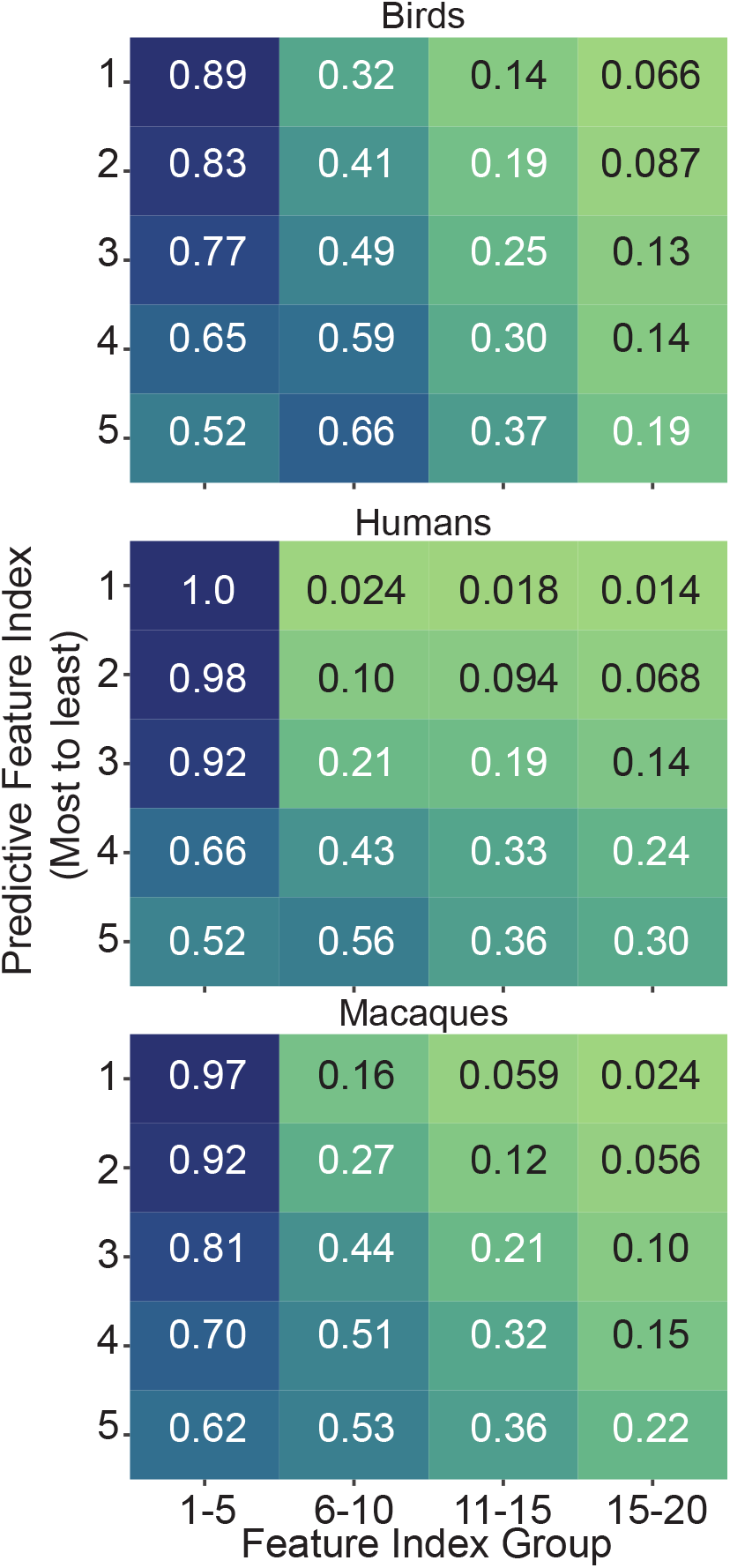
Slown features are predictable features. Canonical Correlation Analysis (CCA) heatmaps (Methods - PFA to SFA Comparison) comparing slow (x-axis) and predictive features (y-axis) for birds, humans and macaques. Smaller indices indicate slower/more predictive features and darker blues indicate higher correlation. Cells show the CCA score (range 0-1 = zero to perfect correlation).

The first five SFA features correlated highly with the three most predictable PFA features (Fig. 7). The fourth and fifth most predictable feature correlated highly with both the SFA features 1-5 and 6-10.

These results show that learning the slow features of vocalizations simultaneously learns their predictable features.

### G. Discriminating vocalizations

We then asked if the “slow” acoustic subspace shared by vocalizations can be used to solve auditory tasks. To address this question, we tested whether a network could discriminate between pairs of conspecific vocalizations using only the features in this subspace.

First, we constructed inter- and intra-class vocalization pairs for each animal group. For birds, intra-class pairs were from the same species while inter-class vocalizations were from different species; for humans intra-class pairs were different or identical digits said by an individual and inter-class pairs were the same digit said by different speakers (Audio MNIST - a standard database for benchmarking audio processing algorithms [43]) ; for macaques we used a similar definition as humans – intra-class pairs were different examples of *coos* from a speaker and interclass pairs were *coos* from different macaques (Methods - Inter- and IntraClass Discrimination Task). We constructed 100 examples of each kind of pair for each animal group (300 inter-class and 300 intra-class pairs in total). Next, we used a simple cochlear model (Methods - Simple Cochlear Model) to generate a cochleagram for each vocalization, concatenated the cochleagrams for each vocalization in a pair, and applied Slow Feature Analysis (SFA).

The time-varying, transient nature of acoustic objects suggests that the auditory system must perform online and dynamic computations [28–30, 44–46]. Indeed, stream segregation, in which mixed inputs are parsed into sources and objects based on attentional goals, is at the heart of auditory perception [44] and seems to require such computations [3, 28, 30, 32, 47–49]. Previous work has shown that SFA can be implemented in an online fashion that dynamically calculates the slowest features of time-varying input signals [21, 37]. Nevertheless, for computational simplicity, we used a batch version of SFA, but to mirror, at least partially, the dynamism of the auditory system and online SFA, we reapplied the algorithm to each vocalization pair individually. Thus, the algorithm approximated the dynamic temporal feature extraction that we expect in audition.

We then projected each vocalization pair onto its slowest SFA features and used this projection to train a simple multilayer perceptron (MLP, Methods - Multilayer Perceptron, Fig. 8A) to discriminate between vocalizations. We started by using the 20 slowest features as we have previously found classification performance using a different variant of SFA maximized at this number of slow features [31]. The classifier achieved high performance for all animal groups and both inter-class and intra-class discrimination. The results were well above chance (Fig. 8B *left* ): Birds Intra-class 99.87% (99.71-99.96%); Humans Inter-class 98.39% (97.40-99.3%); Humans Intra-class 97.70% (96.23-98.87%); Macaques Inter-class 99.94% (99.90-99.96%); Macaques Intra-class: 99.94% (99.90-99.96%) – Median and Interquartile range indicated. The greatest variance in performance occurred for human vocalizations. These results indicate that shared “slow” subspace of animal vocalizations can indeed be used to solve a fundamental auditory task.

**FIG. 8.**
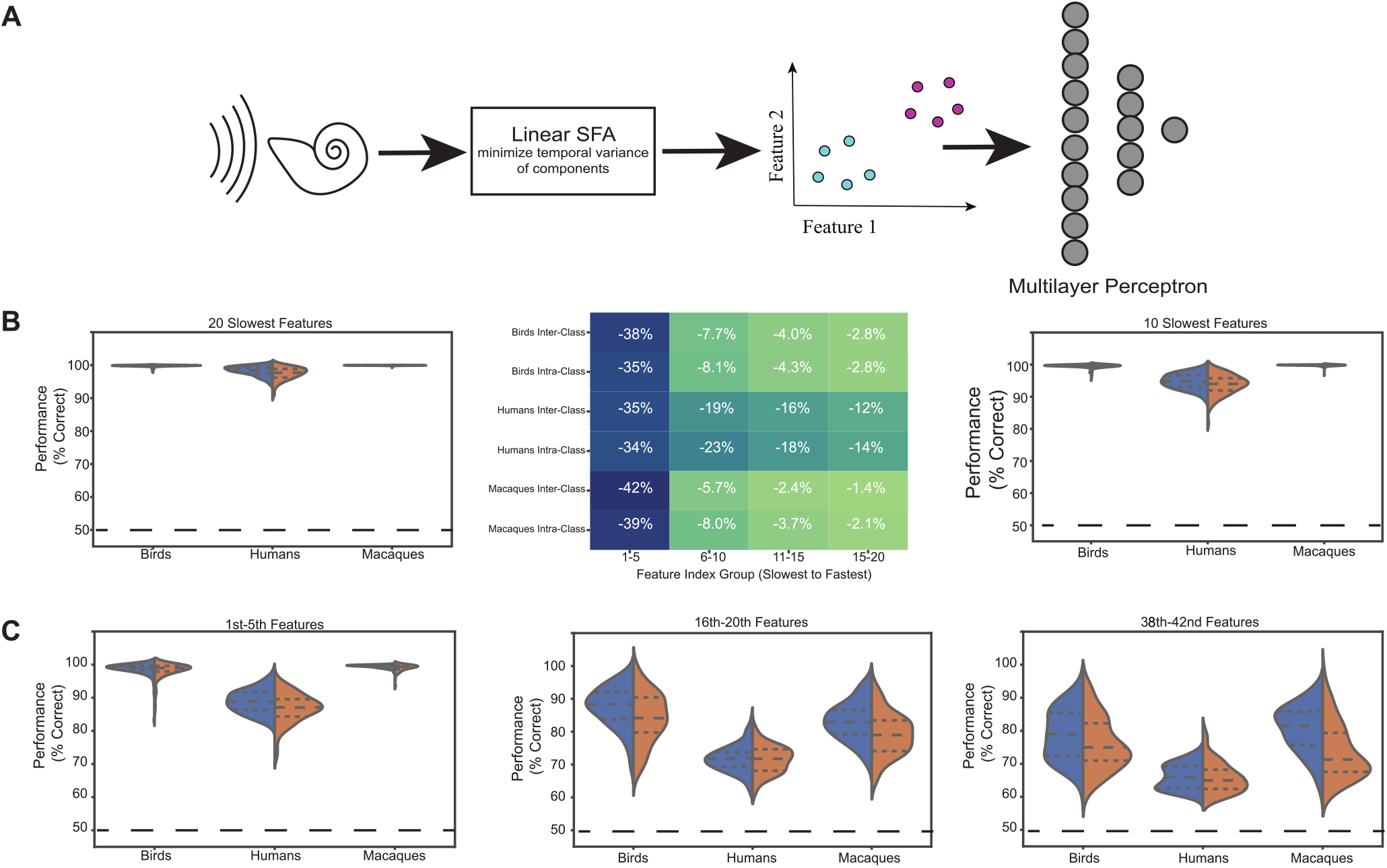
Analysis procedure and discrimination results. **A**. Schematic of analysis pipeline. **B**. *Left:* Violin plots = % correct for inter-class (blue) and intra-class (red) classification with 20 slowest features on 100 vocalization pairs. Birds: intra-class stimuli from the same species; inter-class stimuli from different species. Humans: intra-class stimuli = same or different digits from one speaker; inter-class stimuli = same digit from different speakers. Macaques: intra-class stimuli = different *coo*s from an individual; inter-class stimuli = *coo*s from different individuals. Median (central dashed line) and interquartile range (upper and lower dashed lines) indicated. Chance performance is 50%. *Middle:* Heat map of performance decrease caused by shuffling the feature set on the x-axis before classification. Darker blues = greater performance decrease. Rows = animal group and classification types. *Right:* Same as left panel, but for classification with the 10 slowest features. **C**. Same as left panel of **B** but with subsets of five progressively faster features.

### H. The slowest features are best

Given the results above, we expected the slowest features to play a more important role than faster features for discrimination. To test this, we conducted a shuffling procedure just prior to classification to interfere with the mapping between features and vocalization identity learned by the network during training. We randomly shuffled the feature labels in groups of five from the slowest to the fastest of the first 20 features (I.e., first case: features 1-5 are shuffled, features 6-20 are unaltered).

We then tested the network and measured the percent decrease in classification performance as a function of which subset of features were shuffled. We saw that the five slowest features had the greatest impact on performance followed by the fifth-tenth slowest features. Fig. 8B *Middle* shows the average *decrease* in performance for each subset of features. Human vocalizations show more decrease in performance when any subset of features is shuffled than the other animal groups, consistent with the higher variability in classification performance for this group.

We predicted from our prior SFA inflection point and subspace analysis that a network trained on only the ten slowest features would achieve performance similar to the complete network. Indeed, the reduced feature set obtained similar performance (Fig. 8B *Right* ): Birds Inter-class 99.69% (99.53-99.85%) ; Birds Intra-class 99.65% (99.39-99.89%); Humans Inter-class 94.83% (93.08-96.72%); Humans Intra-class 94.00% (91.19-95.75%); Macaques Inter-class 99.86% (99.78-99.92%); Macaques Intra-class 99.85% (99.73-99.92%) – Median and Interquartile range indicated. The slight increase in variance is expected because we are using fewer features and hence fewer parameters [50, 51].

Our results from shuffling subsets of features led us to ask how well a network trained on only the 5 slowest features would perform. We find that while median performance appears to stay similar, there is a marked increase in performance variability, particularly for humans (Fig. 8C).

Finally, we wondered whether networks trained on any subset of 5 slow features could perform the task. Since we have 42 spectral channels in our cochlear model and SFA is a full spectral technique, we had a max of 42 features to choose from. We chose to look at the 5 fastest features that we used in our initial networks (features 16-20) as well as the 5 fastest features we could find (features 38-42). Performance decreased and variability increased monotonically with feature speed. Interestingly, a network using the five fastest features still had above chance performance. This finding likely relates to previous work showing that task relevant information is often distributed redundantly across sensory or perceptual axes [52].

## III. DISCUSSION

We have used Slow Feature Analysis to reveal a shared and exploitable temporal structure inherent to animal vocalizations. This structure reflects, and also leverages, the concentration of power at low temporal modulation previously reported for vocalizations [18, 19, 27–31]. There is evidence for circuits in the auditory system that tune rapidly to temporal modulation structure in sounds [19, 29, 33, 34, 46, 47]. Such circuits could in principle implement SFA-like computations [21, 31, 37], but their dynamics and response properties have not been fully elucidated. Our results suggest that this would be a fruitful avenue for research. As a first step, one could use regression-based methods such as those in [53–56] to test whether neural activity correlates with the slow features of a stimulus. That is, one could calculate the slow stimulus features, project the stimulus into the SFA feature space, and use the feature magnitudes as regressors to predict firing rates of neurons recorded during stimulus presentation. If the SFA regressors capture significant neural variance, these cells, or perhaps the circuit pathway leading to them, may encode SFA-like features.

We found that about 10 of the slowest features of vocalizations contribute the most to their overlapping structure as well as to vocalization discrimination performance. These results suggest that the slowest SFA features define a shared, low dimensional subspace that is both learnable and useful for auditory tasks. Consider, for example, how this subspace could be used to solve a classic auditory challenge - the cocktail party problem. First, the incoming acoustic stream [44] should be decomposed into its composite frequencies which are then analyzed as a set for something akin to slowness, tracking the 9-11 slowest features. Projecting the signal into this subspace should cause the speaker to “pop out” as their vocal signal will change more slowly than the background noise. Geometrically speaking, the background noise will perform a pseudorandom walk close to the origin of the slow acoustic subspace, while the speaker’s vocal signal will draw a longer, more defined path [30, 31]. Likewise, the more familiar we are with the voice and the longer we listen to it, the more defined the slow features of the speaker’s voice can become, and the more accurately and finely we can parse what they are saying. Tuning to a new speaker requires a rotation of this subspace to align with their slowest features. Since we observed that the slow subspaces of different individuals are related by small rotations, a transformation could be performed in real time and reversed as attention shifts between speakers. It is unlikely that these slow features are learned *de novo* each time. Rather we imagine that a prior is learned based on experience with human voices in general as well as experience with the language spoken specifically. This prior can then be refined online based on the current acoustic context and auditory goal. Presumably there must be some online refinement to account for the changes in the frequency content that can occur between different human speakers [57].

Returning to our initial motivation of parsing conspecific and allospecific vocalizations, the overlap in the slowest features up to small rotations between animal groups likely allows a subspace learned on conspecific vocalizations to still partially work on allospecific vocalizations. At a minimum it seems likely that the overlap would allow the auditory system to identify that the incoming acoustic signal is a vocalization and that it is allospecific or otherwise uncommon. That is, a bird can parse that a cat’s meow is a vocalization not an environmental sound and that it is not a bird vocalization. Conversely, extensive experience with allospecific vocalizations would likely allow for learning good priors on their slow features and thus finer discrimination, as is anecdotally seen with owners recognizing their specific pet’s vocalizations.

In addition to the above regression analyses, one could test the effect of manipulating slow features on perception and behavior to explore to what extent the mechanisms suggested above are implemented in the brain. For example, male cowbirds sometimes sing over competing males and females can produce “chatter” to seemingly block unwanted male songs [6, 58]. Perhaps such chatter calls or singing-over behaviors specifically target slow song features, affecting perception by conspecific individuals. More generally, the slow features of stimuli could be parametrically masked to determine the effect on perception. Many researchers have probed how changes to auditory stimuli affect auditory object perception, but experiments have generally used artificial stimuli due to the challenges associated with systematically parameterizing the temporal statistics of natural stimuli [28, 44, 59–61]. Our results suggest that manipulating slower features of vocalizations, learned through SFA analysis, will have stronger behavioral effects than manipulating faster features, thus providing a systematic basis for design of experiments. Furthermore, our results add to the growing call for ethology and neuroscience to collaborate closely during stimulus and task design [48, 54, 56, 60–70].

On a theoretical level, our results inform the use of frameworks like efficient coding theory to understand the functional organization of neural circuits. The broad idea of efficient coding is that neural circuits are costly and were selected in a manner that produces the greatest benefit for the investment [71]. This approach has explained many aspects of early vision, including nonlinearities in the fly visual system [72], center-surround receptive fields of neurons in the vertebrate retina [73–79], spike timing statistics [80], the preponderance of OFF cells over ON cells [81, 82], the mosaic organization of ganglion cells [83, 84], the scarcity of blue cones and the large variability in numbers of red and green cones in humans [85], selection of predictive information by ganglion cells [86, 87], and the expression of ion channels in insect photoreceptors [88]. Similar analyses suggest that the olfactory periphery [89–91] is also adapted to the statistical structure of the environment to use limited resources efficiently to represent odor information. In visual cortex, adaptation to natural stimulus statistics seems to explain the functional structure of edge detectors in V1 [92], and texture analyzers in V2 [93–95]. Even the grid cells of the entorhinal cortex appear to be efficiently organized to represent space [96]. Some studies have used such ideas [97–100] or related [101–104] to broadly study auditory circuits. Our results suggest that it will be productive to examine in detail how different parts of the auditory pathway may be adapted to efficiently process rich, but low dimensional, temporal regularities of ethologically relevant stimuli.

## IV. METHODS

### A. Auditory Stimuli

We used publicly available vocalization datasets for humans, macaques, and birds (blue jay, house finch, American yellow warbler, and great blue heron). The human vocalizations included six English speakers pronouncing digits zero through five 50 times each (a total of 50^*^6 = 300 pronunciations per person) [43]. Macaque vocalizations were all *coo*s [2, 19, 31]. The data set included 8 individuals with approximately 1000 coos from each individual [105]. Bird vocalizations were not labeled by individual, but were grouped by species, and included approximately 250 song examples from each species [106]. Each animal group was recorded at a different sampling rate (humans at 48 kHz, birds at 44.1 kHz, and macaques at 32 kHz, respectively). We accounted for these differences by including the appropriate sampling frequency in the gammatone filter bank of our cochlear model (see below). We selected vocalizations randomly from these dataset to create vocalization pairs. If vocalizations were accidentally paired with themselves, that pair was eliminated from further analysis. This was rare (occurring *<* 5% of the time) and resulted in rejection of 29 of the 600 total pairs. See Inter- and Intra-Class Discrimination Task below for more details. Due to the large number of vocalizations in each data set, we assumed our examples covered the variance seen in vocalizations, e.g. vocalizations been shorter or longer than average, pitched up or down, higher or lower in volume, etc. We therefore did not artificially augment the data set by performing any manipulations on the original vocalizations as is commonly done in deep learning contexts for more complex, but less biologically plausible, networks [107– 109].

### B. Simple Cochlear Model

We implemented a simple cochlear model to approximate the structure of inputs to the auditory system. The model included two filter stages: 42 gammatone spectral filters followed by a temporal filter following previous work [31, 98, 110, 111]. Each of the 42 filter outputs (42 spectral × 1 temporal filters) was normalized to zero mean and unit variance. The gammatone filters were implemented using the `GammatoneFilterbank` function from python module `pyfilterbank.gammatone`. The gammatone filters had center frequencies between 22.9 Hz to 20.8 kHz, covering the frequency range of all vocalizations studied (approximately 20 Hz to 10 kHz) . The temporal filter was implemented as a difference of two kernels of the form

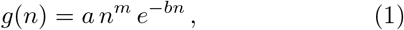

where *n* is in units of samples and *a, b*, and *m* are shape and rate parameters. The temporal filter was created by taking the difference between filters *g*_1_ (*a* = 1.5, *b* = 0.04, and *m* = 2) and *g*_2_ (*a* = 1, *b* = 0.036, and *m* = 2). These parameters mimicked aspects of cochlear temporal processing [111, 112]. Each filter output was normalized to zero mean and unit standard deviation.

### C. Slow Feature Analysis

The conceptual underpinning of Slow Feature Analysis (SFA) is the “slowness principle” [20–22], which hypothesizes that higher-order information in a stimulus (e.g., stimulus identity) changes more slowly than nuisance fluctuations. Linear SFA is defined by the equations

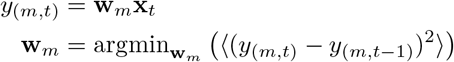

where **x**_*t*_ is an auditory stimulus (or set of stimuli) at each time *t*, **w** is the set of slow features, *y*_(*m,t*)_ is the projection of the stimulus into the feature space defined by **w**, and ⟨⟩ is the expected value. “Slowness” is realized by finding the particular set of features that minimizes the average squared temporal difference of each m component of *y*_*m*_. Because the temporal difference *y*_(*m,t*)_ − *y*_(*m,t−*1)_ is the discrete analog of a derivative, the minimization in the second line selects features to generate a projected output that changes most slowly from moment to moment. More complex versions of SFA replace the first equation with a nonlinear feature extractor.

The optimization problem above is subject to three constraints, which ensure non-trivial solutions:

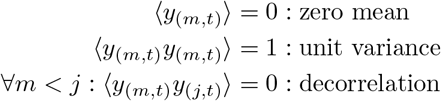

The second and third constraints ensure that the SFA features are linearly independent and order them by slowness, similarly to the ordering induced in principal component analysis (PCA). If the number of features is taken to be smaller than input dimension SFA performs dimensionality reduction.

Also similarly to PCA, SFA can be formulated as a generalized eigenvalue problem. For PCA, we seek eigen-vectors of the data correlation matrix with the largest eigenvalues as these capture the most data variance. For SFA, we instead seek eigenvectors of the temporal difference correlation matrix (i.e., eigenvectors of Δ*X*Δ*X*^*T*^ where Δ*X* = *X*_*t*+1_ − *X*_*t*_) with the *smallest* eigenvalues. These capture directions of least temporal variance, i.e. where the signal changes slowest. For a comparison of PCA and linear SFA see [35].

### D. Eigenvalue Analysis

The Slow Feature Analysis (SFA) optimization problem is often posed as a generalized eigenvalue problem [20–22, 35]. When posed this way, we can interpret each slow feature as an eigenvector with an eigenvalue proportional to its slowness, so that smaller eigenvalues indicate slower features. We can construct distributions of these eigenvalues by performing SFA on many vocalizations, recording their eigenvalues, and averaging across vocalizations. We performed this analysis for each animal group. We calculated the eigenvalues for each intraclass vocalization pair and then averaged across all vocalization for each animal group. A given vocalization can appears multiple times in the calculation, but this was statistically unlikely given the number of vocalizations (see Auditory Stimuli). The calculation generated the three distributions plotted in 4. Since SFA requires that each slow feature be orthogonal to all other features, the distribution is monotonically increasing. That is, each subsequent slow feature will be faster than or as slow as the previous slow features and, equivalently, each eigenvalue will be greater than or equal to the previous eigenvalues. We analyzed the rate of change of slowness across the features by finding the first inflection point on the monotonically increasing eigenvalue distribution of each animal group. This inflection point represents the greatest second derivative along the distribution, after which the eigenvalue, and hence slowness, varies less over the SFA features. We found this inflection point using the KNEEDLE algorithim [38].

### E. The Gram Matrix and subspace overlap

A central claim of this work is that the temporal regularities extracted by SFA are not idiosyncratic to individual vocalizations, but are instead shared within and across animal groups. Establishing this claim requires a quantitative measure of the extent to which the slow features learned from one vocalization agree with those learned from another. Two considerations constrain the choice of such a measure. First, vocalizations differ in duration, amplitude, and spectral content, so the comparison must be made between the learned features (weight matrix *W* ) rather than between the projected time courses (*Y* = *W*^*T*^ *X*), which are not defined on a common time axis. Second, SFA returns an ordered basis for a subspace, and the assignment of structure to individual basis vectors is not unique: two vocalizations may span the same subspace while distributing that structure across their features differently, for instance if the slowest feature of one vocalization corresponds to a mixture of the two slowest features of the other. An adequate measure must therefore be insensitive to rotations within each subspace. We constructed such a measure by calculating the Gram matrix between two learned weight matrices, and then its singular value decomposition (SVD.

The singular values after SVD are invariant to rotations, and characterize the relative orientation of two subspaces provided the feature sets obey two conditions. First, each features must be of unit length, so that an inner product between features is the cosine of the angle between them. Second, the features of a single vocalization must be mutually orthogonal, since the principal angles used below are defined as the arccosines of the singular values of the inner product matrix between orthonormal bases for two subspaces; in the absence of orthogonality the singular values of 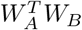 may exceed unity and reflect the conditioning of the chosen bases rather than the geometry of the subspaces themselves. Both conditions are ensured by the manner in which we solve the SFA optimization problem.

For any two vocalizations *A* and *B*, we computed the Gram matrix of their learned weight matrices:

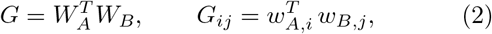

*G* is a 20 × 20 matrix whose (*i, j*) element is the cosine similarity between the *i*th slow feature of *A* and the *j*th slow feature of *B*. Its entries therefore quantify agreement at the level of individual, slowness-ordered features: element (1, 1) compares the slowest features of the two vocalizations. Each entry of *G* refers to a particular pair of features, so the entries change if either set of features is rotated within the subspace it spans, even though the subspaces themselves are unchanged. To compare the subspaces rather than the features, we decomposed *G* as

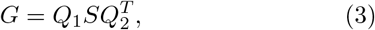

where *Q*_1_ and *Q*_2_ are orthogonal and *S* is diagonal. Because *W*_*A*_ and *W*_*B*_ are orthonormal bases, the singular values *σ*_*i*_ lie between zero and one and are the cosines of the principal (canonical) angles between the two subspaces,

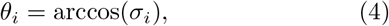

ordered from smallest to largest. These angles describe the relative orientation of two subspaces completely and independently of any rotation within either: an angle near 0^*°*^ identifies a direction shared by both subspaces up to such a rotation, while an angle near 90^*°*^ identifies a direction present in one subspace and absent from the other. Principal angles were computed for every vocalization pair and the resulting distributions averaged within and across animal groups.

### F. PFA to SFA comparison

Predictable Feature Analysis (PFA) [25, 26] seeks the components of a signal that permit best prediction of its future by a pre-specified model. We utilize a linear autoregressive model which is easy to fit and mathematically tractable. We follow the optimization procedure in [26] to find the predictive features, via a generalized eigenvalue problem. The eigenvectors produced by this procedure are the predictive features. While linear SFA generates a set of slow features which can be sorted by slowness, PFA with a linear auto-regressive model generates a set of predictive features which can be sorted by predictability according to the magnitude of each predictable feature’s eigenvalue. We can then compare the slowest and most predictive features.

As discussed in the main text, predictability and slowness are related but not equivalent [25, 26]. A signal could change rapidly but be highly predictable (e.g., a binary signal that changes between 0 to 1 every time step is very fast but also very predictable). Thus we asked whether the predictable components of vocalizations were similar to their slow components. We performed this comparison by using Canonical Correlation Analysis (CCA) [39, 40]. CCA can be viewed as an extension of univariate correlation for comparing multidimensional data sets. There are many variations of CCA, but in its most basic form this method finds the left and right singular vectors of the correlation matrix between two data sets that has the greatest singular value. These singular vectors generate the projections of the two data sets that have the greatest correlation. In our case, the calculation is simpler since we consider each PFA feature individually. When one of the data sets is univariate, CCA reduces to linear regression with the higher dimensionality vectors used as regressors to fit the univariate data. Thus our CCA correlations are effectively the R-squared value when SFA is used to predict the PFA signal. For more details on forms of CCA see [42], and for rigorous deviation of basic CCA see [41]

### G. Inter- and Intra-Class Discrimination Task

To determine whether slow features are sufficient for auditory discrimination, we created a battery of test pairs at random for each animal group. We considered inter-class and intra-class pairs reflecting two broad kinds of discrimination/classification problems: discriminating between different clusters (inter-class) vs discriminating between different examples within a cluster. For humans, we defined inter-class pairs as different speakers saying the same digit and intra-class pairs as the same speaker saying different digits or giving different pronunciations of the same digit. For macaques, we used a similar definition: inter-class pairs were different individuals producing *coo*s while intra-class pairs were different *coo* examples from an individual. For birds, we defined inter-class pairs as vocalizations from different species and intra-class pairs as vocalizations from the same species. Inter- and intra-class vocalizations pairs were analyzed the same way: each member of the pair was passed through the cochlear model to produce a cochleagram, i.e., a matrix showing the power in each frequency group at each time. The cochleagram for each vocalization in the pair was then horizontally concatenated together and analyzed by linear SFA. Next, the vocalization pair was projected into a subset of the feature space found by SFA. This projection results in a cloud of data points corresponding to the SFA projection of the cochleagram at a moment of time. Subsets of these points were then used to train a multilayer perceptron (MLP) to discriminate one vocalization from the other in a pair (see below). The SFA algorithm and the MLP were reset for each vocalization pair.

### H. Multilayer Perceptron

We constructed a simple multilayer perceptron (MLP) to discriminate between vocalization pairs. This MLP consisted of an input layer with as many nodes as the number of SFA features used, a five node hidden layer, a single node output layer. Each hidden layer node used a rectifying linear (ReLu) nonlinearity and the output node used a soft-max nonlinearity. The vocalizations within each pair were randomly assigned the label 0 or 1 and data points belong to that vocalization (cochleagrams frames projected onto the SFA feature space) were labeled accordingly. Thus the output node estimates the probability that input vector belongs to vocalization 1. Inputs with probability greater than 50% were assigned to vocalization 1 otherwise they were assigned to vocalization 0. The network was fit using the ADAM solver [113]. To prevent overfitting, we used five-fold, stratified shuffle split cross-validation with a 50-50 train-to-test split of the data. Briefly, during each repetition the algorithm randomly shuffled the data and sampled 50% of the data from the vocalization pair making sure to maintain the original ratio of data between the two vocalizations. We then sub-sampled the data to ensure that there were equally numbers of data points in the training set. These samples were used for training and and the network was tested on the remaining data, which we also subs-sampled. Sub-sampling ensures that chance performance is 50% rather than the 1-the marginal probability of the shorter vocalizations. This procedure was repeated five times and samples were taken with replacement (i.e., the same sample could appear in multiple repetitions). See the documentation for `StratifiedShuffleSplit` in the `model selection` module of `sklearn` for more details. The average score across the repetitions was reported. We used percent of test data classified correctly as our performance metric.

### I. Programming languages and code availability

Code was written in Python 3.8.19 and will be/is available via the Balsubramanian lab `Github` repository: https://github.com/BalasubramanianLab-CNI. Links to necessary modules are included in the `ReadMe`, or are provided in the repository.

## Acknowledgments

VB and RWD were supported in part by NIH grant R01EB026945 *Coincidence and continuity: uncovering the neural basis of auditory object perception*. LW was supported in part by the Velay Fellowship from the University of Pennsylvania. VB thanks the Aspen Center for Physics (NSF grant PHY-2210452) for hospitality as this work was completed. VB was supported in part by the Eastman Professorship at Balliol College, University of Oxford while this work was being performed.

